# In vitro study of dentin hypersensitivity treated by whitlockite glass-ceramics

**DOI:** 10.1101/226068

**Authors:** Amanda C. Juraski

## Abstract

The aim of this study is to evaluate the potential of a bioactive glass based on the 3CaO.P_2_O_5_-SiO_2_MgO-system and its glass-ceramics containing whitlockite on the remineralization of dentin as a possible treatment to dentin hypersensitivity. For that, 40 third molar human teeth were artificially demineralized and randomly distributed in 4 groups (n = 10): G1 - Negative Control (no treatment), G2 - Positive Control (treated by Bioglass® 45S5), G3 – BG (treated by bioactive glass based on 3CaO.P_2_O_5_-SiO_2_-MgO-system), and G4 – BGC (treated by bioactive whitlockite glass-ceramics). After treatment, the samples were emerged in artificial saliva and stored for 7 days in a controlled temperature of 37ºC. After that, scanning electron microscopy (SEM) and atomic force microscopy (AFM) was used to evaluate samples morphology. The analysis confirmed the formation of hydroxyapatite on the surface of all the biomaterials studied, that in the dentine specimens treated by bioactive glass and whitlockite glas-ceramic most of the dentinal tubules were completely occluded.

## 1. Introduction

Dentin is a tubular, permeable, mineralized structure that composes the most part of the human teeth. This tissue is mainly formed by type I collagen fibrils, glycosaminoglycans, phosphoproteins, phospholipids and hydroxyapatite crystals.

The dentin demineralization can occur due to exposition of this tissue to acids that can be present in the oral environment, as well as a result of abrasion or attrition of dentinal exposed surface. This process is characterized by the loss of hydroxyapatite crystals and, as a consequence, the opening and exposition of dentinal tubules, which can result in hypersensitivity. (Wang et al. 2011)

Bioactive glasses and glass-ceramics are promising biomaterials in the field of bone regeneration and dentin remineralization due to their exceptional bioreactive properties that allows to easily form hydroxycarbonate apatite (HCA) when immersed in simulated body fluids. (Wang et al, 2011).

The concept behind this project is to study the potential of a bioactive glass based on the 3CaO.P2O5-SiO2-MgO system and its glas-ceramics containing whitlockite on the reconstruction of the mineral phase of the human dentin. A qualitative analysis Scanning Electron Microscopy (SEM) and by Atomic Force Microscopy (AFM) was performed to observe the occlusion of the dentin tubules by the formation of hydroxyapatite.

## 2. Materials & Methods

### 2.1 Sample Preparation

Forty extracted sound third molar human teeth were obtained from the Human Teeth Bank of School of Dentistry of University of Sao Paulo. From each tooth it was obtained a 3 × 3 × 1mm dentin slab using a water cooled diamond saw (EXTEC Labcut 1010 Low Speed Diamond Saw). The cuts were made 3 mm below the occlusal surface of the teeth. A standard smear layer was created using a 600-grit silicon carbide for 60 s under constant water irrigation.

### 2.2 Experimental Procedures

Before receiving the biomaterials, the surface of samples was demineralized with a 0.5 M EDTA solution (pH 7.4) during 5 min and then washed with distilled water for 2 min to assure complete removal of the solution. After that, the samples were randomly distributed in four experimental groups (n = 10): G1 – Negative control (no treatment), G2 – Positive control (treated with Bioglass® 45S5), G3 – Treated with bioactive glass from the 3CaO.P_2_O_5_-SiO_2_-MgO system, and G4 – Treated with whitlockite bioactive glass-ceramic from the 3CaO.P_2_O_5_-SiO_2_-MgO system. After treatment, all samples were stored in 20 mL of artificial saliva for 7 days at controlled temperature of 37ºC to induce the remineralization process.

### 2.3 Morphological Analysis

The morphological analysis of the samples was made by using a Scanning Electron Microscope, Phenon/FEI. Atomic Force Microscope, AFM/SPM Series 5500 Agilent, was used to determine the topography of the samples. Each sample had six 70 ×70 µm images and six 40 × 40 µm images taken.

## 3. Results & Discussion

Figure 1A and 1B show the SEM images obtained from G1 and G2 groups. In Figure 1A, it is possible to observe the dentin tissue with the exposition of dentinal tubules, which were fully or partially opened due to the demineralization process. After the application of the Bioglass® 45S5, it was observed the presence of agglomerated material on the dentin surface, with the presence of globules and dentin tubules totally covered. Figures 1C and 1D show the SEM images obtained for samples from G3 and G4.

**Figure 1.**
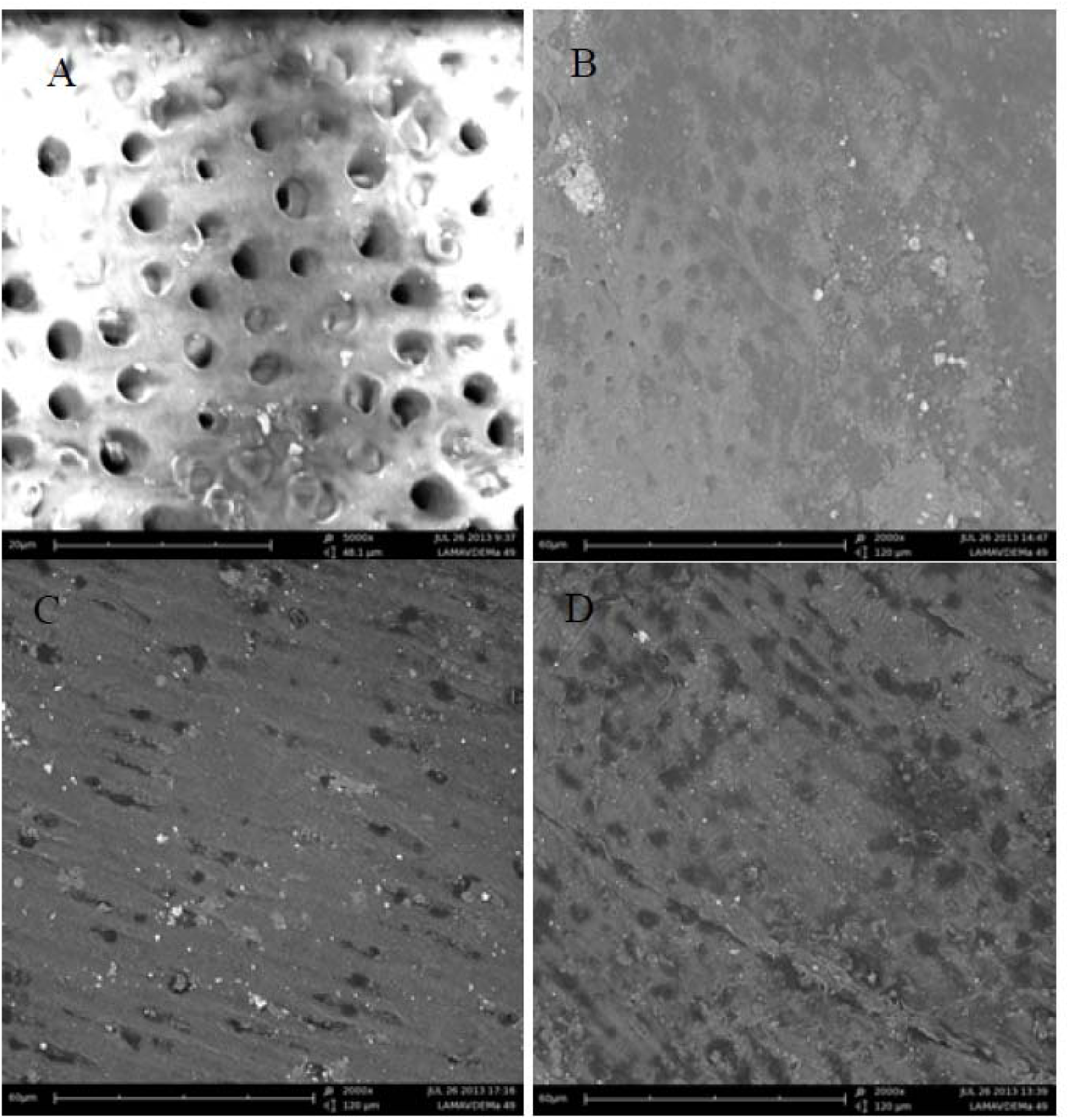
SEM images of samples from: A) G1 - negative control group, B) G2 positive control group, C) G3 – treated with bioactive glass and D) G4 – treated with bioactive whitlockite glass-ceramic. After the process of remineralization. Original magnification: 5000x.

The samples that received the treatment with the bioactive whitlockite glass-ceramic (Fig. 1D) presented a thick layer of hydroxyapatite on the surface, covering completely the tubules that were opened by demineralization. The samples from G3, treated with bioactive glass, also presented their tubules coated by hydroxyapatite.

Comparing figures 1C and 1D it is perceptible that the formation of hydroxyapatite on the samples from G4 was more uniform that on the samples from G3. In Fig. 1D it is possible to notice a nearly continuous precipitate throughout the entire surface of the samples treated with the bioactive whitlockite glass-ceramic, while Fig. 1C shows that, although there is satisfactory layer of hydroxyapatite formed, it does not extend as uniformly throughout the sample surface. Even so, it is clear that the dentin tubules in the samples treated with the bioactive glass and the bioactive glass-ceramic from the 3CaO.P_2_O_5_-SiO_2_-MgO system are completely covered.

The topographic AFM analysis resulted in several 40 ×40 µm images of dentin samples with different treatments. Figure 2A to 2D show the AFM images from G1, G2, G3 and G4, respectively.

**Figure 2.**
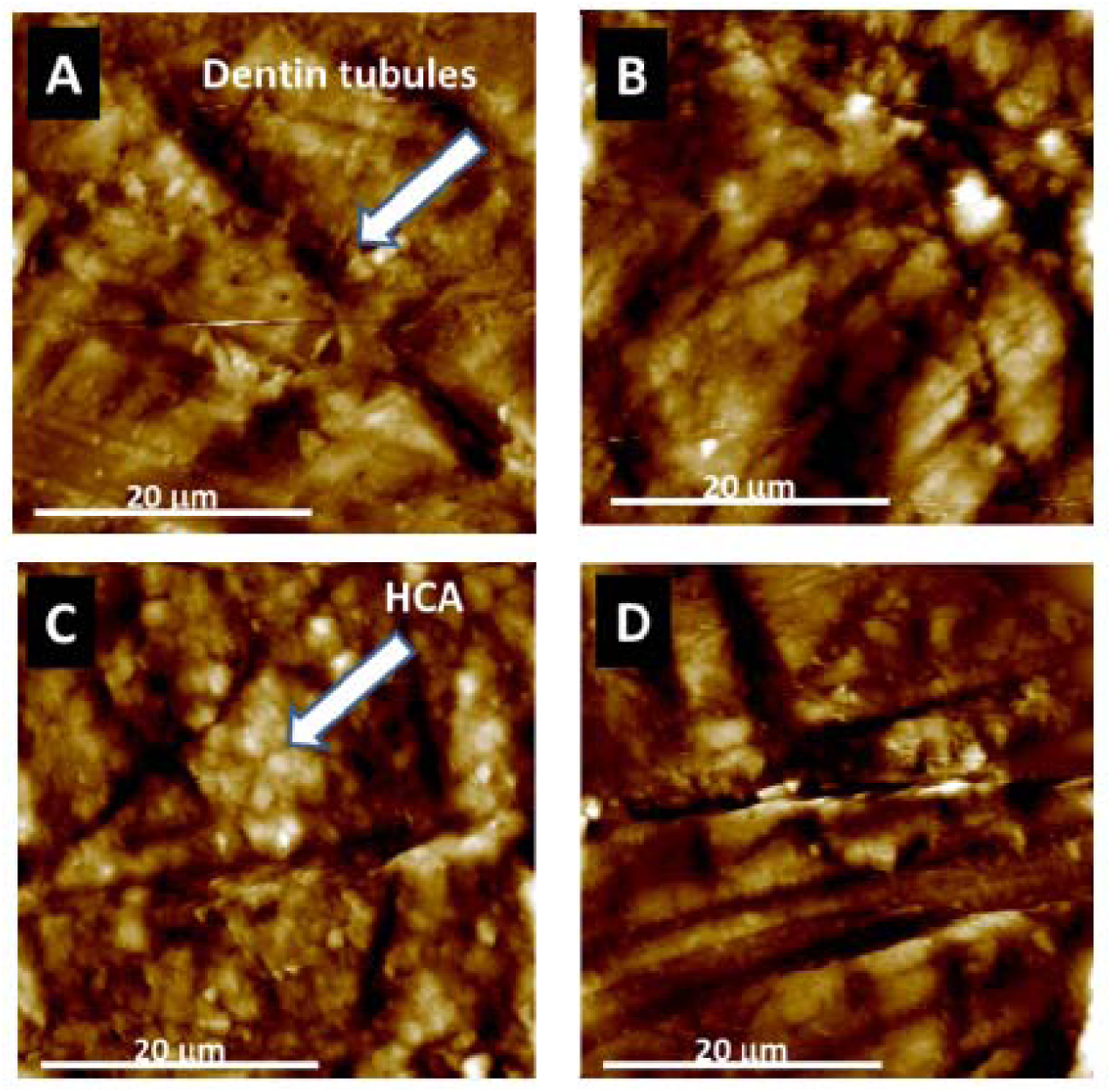
AFM images: A) G1 - negative control group, B) G2 - positive control group, C) G3– treated with bioactive glass and D) G4 – treated with bioactive whitlockite glass-ceramic.

In Fig. 2A is possible to see opened dentin tubules after the seven days immersion in artificial saliva. In the following images (Fig. 2B to 2D) a different type of topology is observed, with the dentin tubules partially covered by the deposition of hydroxyapatite. The images also suggest a smoother surface for the samples that received the some sort of treatment.

With the SEM and AFM images it was possible to observe the morphological and topographic changes that occurred after the remineralization process. The SEM micrographs show that the hydroxyapatite formed on the G4 samples has a more crystalline aspect, due to the presence of whitlockite in the bioactive glass-ceramic (Daguano et al, 2013).

With the AFM images it was possible to observe the morphological and topographic changes that occurred after the remineralization process. The AFM images indicate the occlusion of the dentin tubules by the formation of hydroxyapatite on the surface of the samples. The images can also suggest that the samples that received treatment with a bioactive glass or glass-ceramic have a smoother surface than the sample from the negative control group. A quantitative analysis of the roughness will be necessary to determine the differences at the submicron scale in the topographic features of the surface samples in the several experimental groups (Wang, 2011).

## 4. Conclusion

It was possible to conclude that both the bioactive glass and the bioactive glass ceramic from the 3CaO.P_2_O_5_-SiO_2_-MgO system promoted the formation of hydroxycarbonate apatite on the demineralized dentin, in a way similar to that of Bioglass® 45S5. In this way, both biomaterials have a high potential for a future clinical application for dentin remineralization. Both the bioactive glass and the bioactive glass-ceramic from the 3CaO.P_2_O_5_-SiO_2_-MgO system promoted the formation of hydroxycarbonate apatite on the demineralized dentin, in a way similar to that of Bioglass® 45S5.

